# First Whole-Body Three-Dimensional Tomographic Imaging of Alpha Particle Emitting Radium-223

**DOI:** 10.1101/414649

**Authors:** Diane S. Abou, Andrew Rittenbach, Ryan E. Tomlinson, Paige A. Finley, Benjamin Tsui, Brian W. Simons, David Ulmert, Ryan C. Riddle, Daniel LJ Thorek

**Affiliations:** Department of Radiology, Washington University School of Medicine, Saint Louis, Missouri, USA; LinQuest Corporation, Los Angeles, California, USA; Department of Orthopaedic Surgery, Thomas Jefferson University, Philadelphia, Pennsylvania, USA; Department of Radiology and Radiological Sciences, Johns Hopkins University School of Medicine, Baltimore, Maryland, USA; Center for Comparative Medicine, Baylor College of Medicine, Houston, Texas, USA; Department of Molecular and Medical Pharmacology, University of California Los Angeles; Division of Urological Research, Department of Clinical Sciences, Lünd University, Malmö, Sweden; Department of Orthopaedic Surgery, Johns Hopkins University School of Medicine, Baltimore, Maryland, USA; Department of Biomedical Engineering, Washington University, Saint Louis, Missouri, USA

**Keywords:** theranostics, pharmacokinetics, molecular imaging, alpha particle radiotherapy

## Abstract

Objective: Dose optimization and pharmacokinetic evaluation of alpha emitting Radium-223 dichloride (^223^RaCl_2_) by planar gamma camera or single photon emission computed tomographic (SPECT) imaging are hampered by the low photon abundance and injection activities. Here, we demonstrate SPECT of ^223^Ra using phantoms and small animal in vivo models. Methods: Line phantoms and mice bearing ^223^Ra were imaged using a next generation dedicated small animal SPECT by detecting the low energy photon emissions from ^223^Ra. Localization of the therapeutic agent was verified by whole body and whole limb autoradiography and its effect determined by immunofluorescence. Results: A state-of-the-art commercial small animal SPECT system equipped with a highly sensitive collimator enables collection of sufficient counts for three-dimensional reconstruction. Line sources of ^223^Ra in both air and in a water scattering phantom gave linear response functions with provide full-width-at-half-maximum of 1.45 mm. Early and late phase imaging of the pharmacokinetics of the radiopharmaceutical were captured. Uptake at sites of active bone remodeling were correlated with DNA damage from the alpha particle emissions. Conclusions: This work demonstrates the capability to noninvasively define the distribution of ^223^Ra, a recently approved alpha emitting radionuclide. This approach allows quantitative assessment of ^223^Ra distribution and may provide radiation dose optimization strategies to improve therapeutic response and ultimately to enable personalized treatment planning.

## Introduction

Prostate cancer afflicts nearly 1 in 7 men globally (*1*). When detected early and confined to the organ, it can be successfully treated with external beam radiotherapy or surgery. However, there is no effective long-term treatment for non-organ confined and metastatic prostate cancer. Disseminated disease is often treated primarily with hormonal therapy, to which prostate cancer inevitably develops resistance. This fatal stage of the disease is characterized by metastatic seeding of the axial and then appendicular skeleton. Osseous metastases are often painful, reduce bone quality, reduce structural integrity leading to fracture, and invade the bone marrow and its hematological and stromal compartments. Bone metastases contribute to significant disability and eventually death (*2,3*).

Radium-223 dichloride (^223^RaCl_2_) is a recently approved alpha particle emitting radiopharmaceutical for application in men with castration-resistant metastatic prostate cancer. The radiotherapy agent demonstrated an improved median survival benefit of approximately 14 weeks versus placebo in a large-scale Phase III clinical trial (*4*). Subsequent subgroup analysis endorse these results, even for patients pretreated with chemotherapy. The alpha particle endoradiotherapy approach also provided palliative relief, extended time to first symptomatic skeletal event and reduced incidence of spinal cord compression (*5,6*).

Radium is a divalent cation that is thought to be incorporated within the bone matrix at sites of active remodeling as a calcium mimetic. The majority of the 28.2 MeV emitted by ^223^Ra and its decay chain is in the form of four alpha particles. These charged helium nuclei exhibit high linear energy transfer, damaging nearby cells while sparing distant tissues from ionizing radiation.

The efficacy of the treatment combined with its favorable toxicity profile has encouraged the investigation of strategies towards improved and extended dosing (ClinicalTrials.gov identifier: NCT02023697 and NCT01934790). Currently, ^223^RaCl_2_ is administered on a per weight basis independent of patient disease characteristics or specific uptake distributions. To facilitate improved and personalized treatment regimes and to predict response, several imaging initiatives have been undertaken. Research has focused upon imaging of either surrogate bone-scan agents (such as ^99m^Tc-methyl diphosphonate or [^18^F]-NaF) or direct ^223^Ra emissions to plan therapy (*7,8*).

#### Significance

Radium-223 is the first approved alpha particle emitter and is an effective therapeutic in castrate resistant prostate cancer patients with primarily bone metastatic disease. However, the mechanism of action and an understanding of factors that affect ^223^Ra distribution and its effects are poorly understood. Here, we utilized high sensitivity and high resolution SPECT imaging to quantify the distribution of ^223^Ra. Our work establishes noninvasive molecular imaging as a tool to elucidate the radiobiological effects of this radiotherapy and to optimize its use.

Direct determination of uptake at sites of disease using ^223^Ra itself has thus far been limited to scintigraphic imaging in man (*9*). These efforts are confined to planar scans, as count rates have been deemed insufficient for single photon emission computed tomography (SPECT) reconstruction. Efforts to quantitate images are in the early stages, but are restricted by low count rate, significant scatter, and attenuation (*10-12*). Despite the difficulties, these studies have revealed important features of the *in vivo* fate of the radionuclide. Notably, a considerable proportion of the initial activity is rapidly incorporated into the gastrointestinal tract and excreted. Adverse events caused by dose adsorbed here include intractable nausea and gastrointestinal symptoms (diarrhea, constipation and loss of appetite), which may lead to cessation of treatment (*13*).

We have recently shown that the pharmacokinetics of ^223^Ra in animal models recapitulates that those found in man using *ex vivo* measures including gamma (γ)-counting, alpha camera imaging and whole body autoradiography (*14*). This establishes murine models as an attractive platform to better understand the effects of and to optimize treatment with ^223^Ra. Exciting developments in the SPECT field with respect to both instrumentation and reconstruction have motivated development of next-generation preclinical systems. High-resolution and sensitivity of imaging and therapeutic radionuclides in preclinical models have yielded insight into experimental agent development down to resolution as low as 0.25 mm (*15-17*). Here we report the application of tomographic imaging of Radium-223 in mice for non-invasive delineation of radionuclide distribution at sub-microcurie injection activities with the potential to monitor dose distribution in real time.

## Results

### Characterizing Imaging Properties of Radium-223

The gamma ray spectra of Radium-223 and daughter radionuclides (including Radon-219, Lead-211 and Bismuth-211) were measured using a high resolution HPGe, as shown in Figure 1A. While less than 2% of the energy of the emitted energy of ^223^Ra and daughters is in the form of photons, it can be seen that there are characteristic emissions present for *in vivo* imaging. The NaI(Tl) detectors used in the camera of the small animal U-SPECT system are of significantly lower energy resolution than the cryo-cooled HPGe system. To determine if the U-SPECT could be used to detect and quantify low (biologically relevant) levels of ^223^Ra, a glass capillary tube of 22 kBq (0.59 μCi) of activity was evaluated. A representative, normalized spectra of the rod phantom from one of the three γ-camera heads from a 10 min acquisition is shown (Fig. 1B). As expected, individual emissions are no longer discernable, however the prominent ^223^Ra γ rays are apparent.

**Figure 1.**
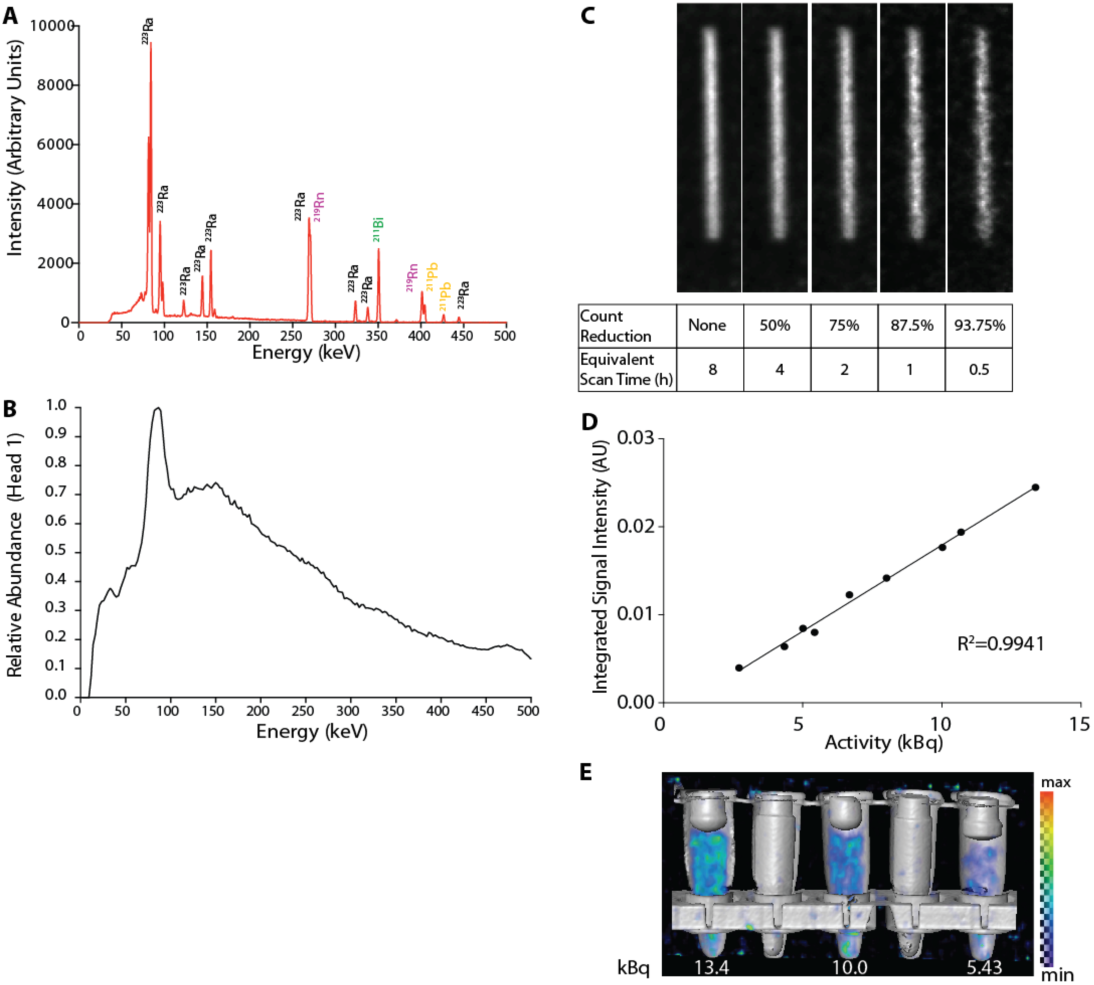
A) Gamma spectrum of Radium-223 and daughters acquired using a high purity germanium detector. The most abundant emissions are the ^223^Ra emissions in the range of 80-100 keV. B) The resolution of the sodium iodide γ-camera is significantly lower than the HPGe, while still detecting the ^223^Ra peaks nearing 100 keV. C) Images of three-dimensional reconstructed line phantoms of ^223^Ra (using an energy window of 80-100 keV) with decreasing count reduction parameters. D,E) Linear response of ^223^Ra activity and user defined volumes of interest, and representative fusion SPECT/CT image, respectively.

### Phantom Imaging of Radium-223 Sources

A line source with 23.5 kBq ^223^Ra was imaged to determine if sufficient counts could be provided for SPECT reconstruction. A list-mode acquisition of 480 minutes was performed using the 80-100 keV energy window. Different fractions of the list-mode data with different totals of acquired counts were used in reconstruction to evaluate the effects of low counting statistics. We found that the small animal SPECT system provided high photon detection sensitivity to provide sufficient detected counts in a reasonable acquisition time for image reconstruction (Fig. 1C).

To evaluate the quantitative accuracy of the SPECT system, samples were imaged following a system calibration with a NIST-standardized ^223^Ra source. Solutions of ^223^RaCl_2_ dissolved in 0.03 M citrate buffer in 0.2 mL Eppendorf microcentrifuge tubes were imaged in the scanner. Volumes of interest of the different activity levels were manually drawn in the reconstructed datasets, and plotted relative to the known activity (Fig. 1D). Representative fusion of the SPECT and CT volumes of the tubes are shown in Fig. 1E.

### Tissue Phantom Imaging of Radium-223

The photon emissions from ^223^Ra used in SPECT imaging are in the range of energies that can be scattered heavily when transported through a biological medium. To determine the effect of scattering material on the system line response function, we imaged a line source in air and embedded in 99% water hydrogel approximating the diameter of a 25 g mouse (Fig. 2A-C). Line intensity profiles across the 0.5 mm inner diameter capillary rod varied only slightly, with a full width half max (FWHM) value of 1.21 mm and 1.31 mm in the vertical and horizontal profile in air and 1.28 mm and 1.41 mm in gel. This compares favorably with the vendor provided reconstructed resolution of <1.1 for this collimator using the 140.5 keV emitting ^99m^Tc.

**Figure 2.**
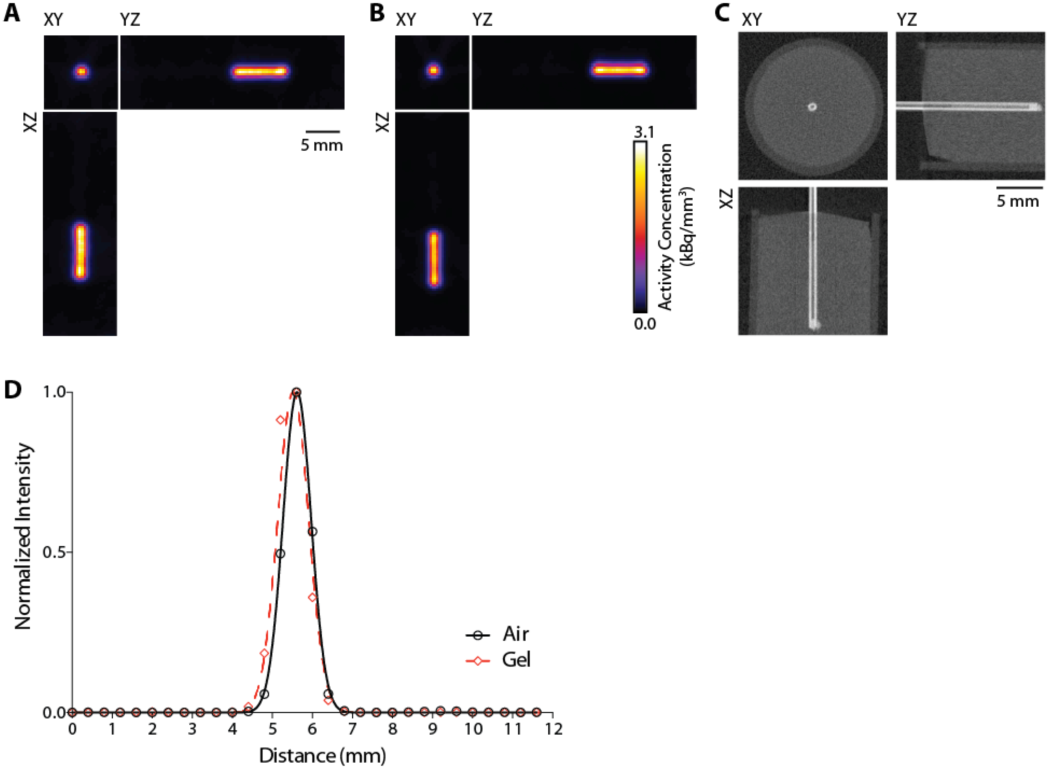
Axial, coronal and sagittal slices through the reconstructed rod phantom in A) air and B) embedded in a 99% water hydrogel to simulate scattering from tissue. Volumes were both reconstructed with 8 subsets and 10 iterations using an iterative pixel based OSEM algorithm. C) CT of the 1 mm external diameter rod source embedded in the hydrogel. Scale bar for all images is 5 mm. D) Intensity profile in air (black) and gel (red dashed) of the horizontal profile across the capillary tube.

### *In vivo* Theranostics of Radium-223

To investigate the *in vivo* distribution of ^223^Ra, we first imaged mice after blood pool clearance and uptake at sites of active bone remodeling (Fig. 3). Animals (n=4) were administered between 20.3-26 kBq (0.55-0.7 μCi) by retro-orbital injection and imaged 24 hours later, when the activity had cleared the soft tissues (*14*). As can be seen in a representative subject, SPECT, CT and fusion images reveal intense localization to the ends of the sites with the highest rates of bone formation and remodeling. This was demonstrated most clearly in the trabecular bone compartment of the distal tibia and proximal femur (Fig. 3A-C). Additionally, signal from ^223^Ra is apparent in the mandible and premaxilla as well as the proximal humerus and distal forelimb. Reconstructed SPECT volumes overlaid with CT reveal the capacity for whole-body tomographic imaging of uptake at sites of interest throughout the skeleton (Fig. 3D,E). Furthermore, as expected, the imaged distribution closely matches that seen using ^99m^Tc medronic acid, the commonly used radiolabeled bisphosphonate bone scan tracer (Supplemental Fig. 1).

**Figure 3.**
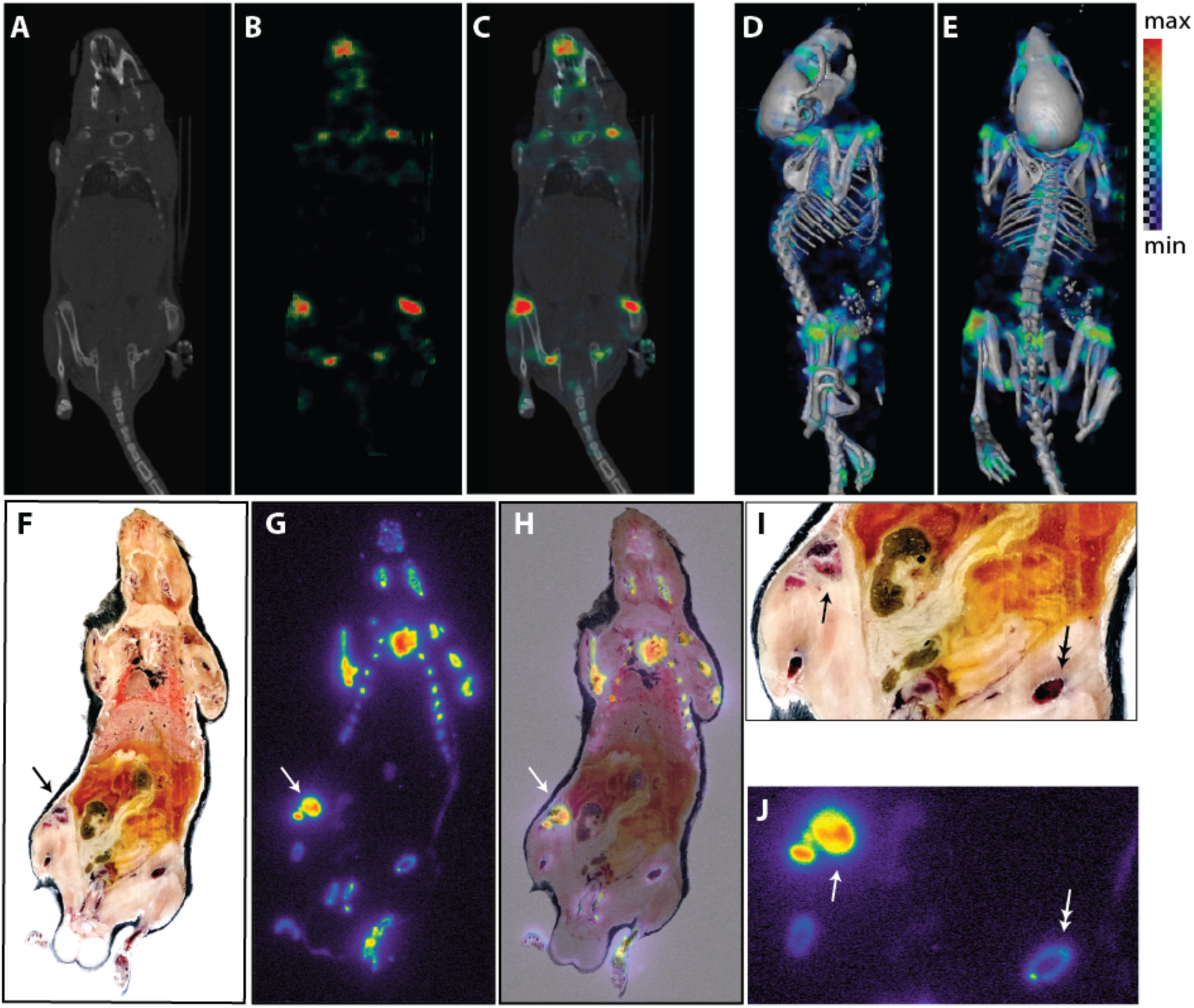
Radium-223 SPECT/CT and Whole Body Autoradiography at 24 h post-injection. A) CT, B) SPECT and C) fused coronal slice of a mouse at 5 min post-injection of ^223^RaCl_2_. D,E) Lateral and dorsal views of three dimensional volume rendered SPECT data overlaid onto surface rendered CT data. To generate this image, a weighted average of the intensities along projections through the SPECT was blended with the CT rendered image. Both slice and three dimensional rendered volumes reveal intense uptake at the ends of the long bones and premaxilla. To confirm *in vivo* SPECT results, whole-body undecalcified cryosections (14 μm) were obtained after imaging. F) En bloc color macrograph, G) autoradiography and H) overlay. I,J) Magnified images of the uptake at the midsection. Arrow indicates the right knee; double-headed arrow the ^223^Ra uptake along the femoral shaft.

### *Ex vivo* Correlation of Radionuclide Distribution and Radiobiological Effect

To verify the noninvasive SPECT images we performed unfixed, undecalcified whole mount whole-body cryo-sectioning and autoradiography. A color macrograph of the animal was acquired en bloc before sectioning (14 μm thickness) and transfer to a storage phosphor sheet for autoradiography (Fig. 3F-H). Consistent with reconstructed SPECT volumes, the overlay image clearly recapitulates the accumulated ^223^Ra throughout the skeleton, notably at the manubrium of the ribcage, lower mandible and hind limbs. Magnified images of the midsection are shown, at the right knee and left femoral midshaft (Fig. 3I,J).

Closer analysis of the leg was performed to assess correlation of imaging results and radiotherapeutic effect. SPECT/CT imaging of the lower hind-limb shows intense uptake at proximal tibia and distal femur at 24 h (Fig. 4AC). The leg was resected a day after the imaging study and en bloc imaging, autoradiography and immunohistochemistry were performed (Supplemental Fig. 2). Labeling was most evident in the rapidly mineralizing bone of the metaphysis. We subsequently evaluated therapeutic effect by annotating DNA damage with immunofluorescence microscopy in treated and control animals (Fig. 4D,E). Sites of phosphorylated γH2AX were found throughout the trabecular bone compartment of the proximal femur in the treated animals in comparison to the low background staining in untreated control samples.

**Figure 4.**
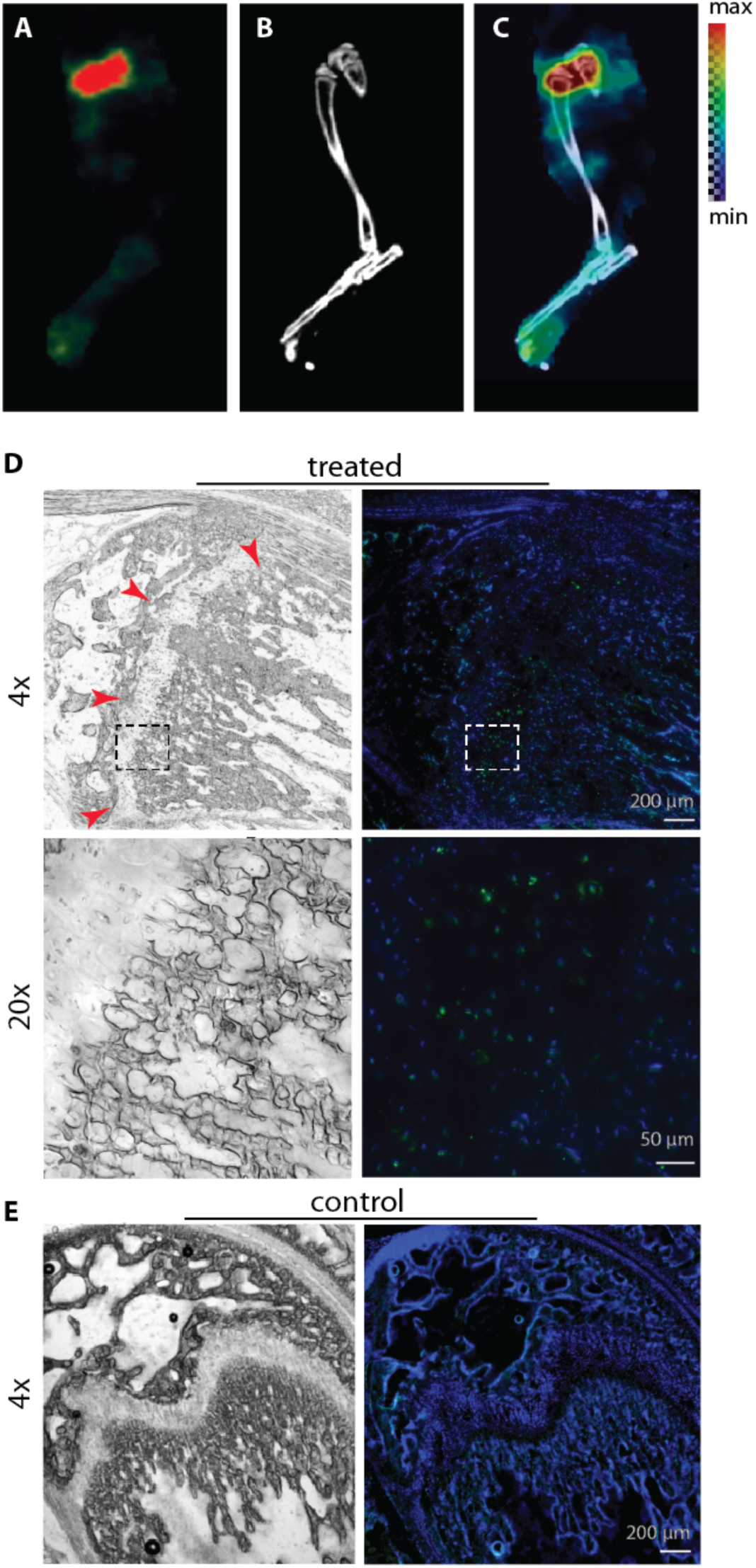
A) SPECT, B) CT and C) fusion of ^223^Ra in the lower hind limb. Uptake is seen at the ends of the long bones where bone turnover and ossification are localized. White light (left) and phosphorylated γH2AX immunofluorescence staining (right) of the femoral head in D) ^223^Ra-treated and E) control mice. Foci of phosphorylated γH2AX staining are identified throughout the bone forming front of the treated animals (red arrows).

### Imaging of Transient Radiotherapy

Our attempts at kinetic imaging of the low injected activities of ^223^Ra were not successful due to extremely low detected counts. However, static imaging of subjects at defined intervals after injection can be used to approximate distribution at arbitrary time points. As an example, 2 minutes following administration of a 16 kBq (0.43 μCi) dose, the animal was sacrificed, and imaged (Fig. 5A-E). At this early time point, the agent is detected in the heart. After SPECT/CT, the animal was rapidly cryo-sectioned and processed for autoradiography (Fig. 5F-H). Pooled activity in the heart and vena cava are identified, in addition to incomplete labeling of the skull.

**Figure 5.**
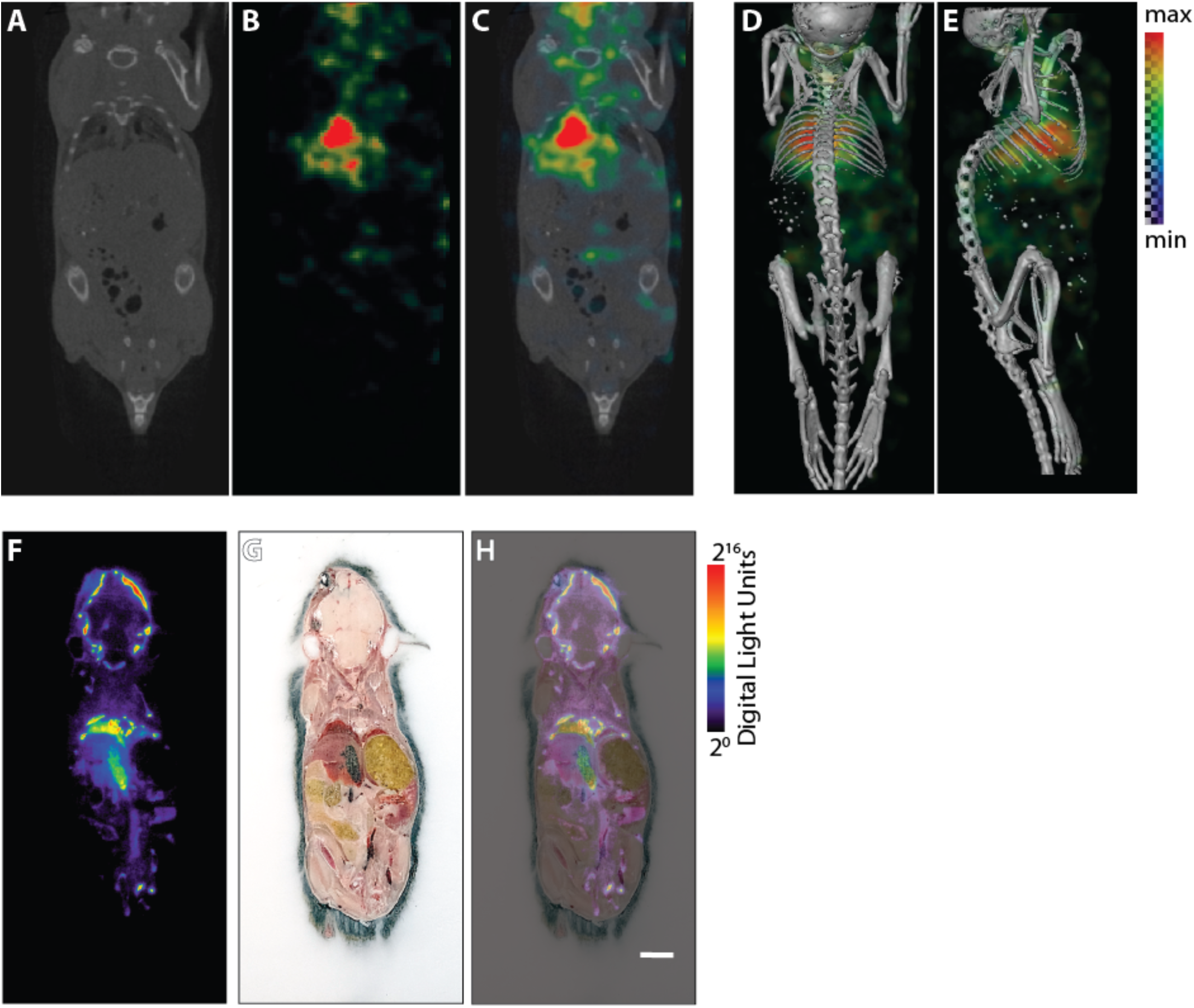
Radium-223 SPECT/CT and Whole Body Autoradiography Immediately After Administration. *In vivo* A) CT, B) SPECT and C) fused coronal slice of a mouse sacrificed 2 min post-injection of ^223^RaCl_2_. D) Tomographic SPECT/CT fusions at dorsal-ventral and E) lateral projections. F-H) Autoradiogram of cryosectioned whole animal, en bloc macrograph and fusion.

## Discussion

The increasing interest in the use of alpha particle emitting radionuclides for cancer cell therapy and ablation is an exciting development in the field of nuclear medicine. A key element in realizing the potential of this emerging strategy is to refine dosing parameters for maximized therapeutic effect. Tomographic imaging can play a significant role in achieving improved pharmacokinetic modeling and dose optimization of radionuclide tracers and therapeutic compounds but has largely not been possible for alpha particle emitting radionuclides.

To date, scintigraphic imaging of ^223^Ra (*8-11*) and antibody-labeled ^213^Bi (*23*) have greatly informed upon the biodistribution and estimated absorbed dose in several patient studies. The primary advantage of a tomographic SPECT is to remove out-of-plane information, rather than the planar images consisting of projections of the 3D activity distribution. SPECT provides improved contrast, resolution and the capacity for true quantitation. Here, we evaluated the imaging characteristics and *in vivo* imaging capabilities of a dedicated small animal SPECT system possessing high sensitivity (*16*). This work demonstrates that the acquisition of non-invasive, quantitative, tomographic distribution of Radium-223 can be achieved using a system with high sensitivity.

An energy window from 80-100 keV was used to acquire three-dimensional distribution of submicrocurie activity levels of ^223^Ra through the detection of the low level of photon emissions. Phantom studies demonstrate that near millimeter resolution and scan times equal to other preclinical nuclear medicine procedures are attainable. The linear response of activity in the investigated region can be used to determine activity concentrations non-invasively. As expected, ^223^Ra labels the sites of active bone remodeling throughout the skeletal compartment. The non-invasive *in vivo* imaging performed well to highlight areas of intense uptake, such as at the mineralizing front of the metaphysis in the long bones, without the onerous requirements of fresh-frozen cryo-sectioning and autoradiography. The cytotoxic effects of the four alpha particle emitting ^223^Ra require further study to optimize activity amounts and scheduling in patients. We have shown that imaging correlates with immunofluorescent localization of DNA damage, which may inform upon these questions.

Limitations of this work include the low signal-to-background and the limited field of view of the system. The former can be overcome with higher injected activity, amounts which may also enable more rapid dynamic imaging. While such doses may indeed be tolerated with only mild toxicity (*24*), they severely depart from clinical practice. Furthermore, background readings made from the system indicate the non-ideal shielding of the γ-camera detectors of the SPECT in our animal facility (containing other photon-emitting radioactive animals within several meters’ distance). With improved facilities management, one could expect improved signal-to-noise.

The second issue, relating to the small field of view, is a system dependent parameter. The imageable volume consists of a large number of projection angles at high magnification focused within the collimator, and whole body coverage is provided by bed motion through the detector. Redesigned collimators may enable the functional imaging of larger subjects, such as mature rodents of interest for research study. ^213^Bi sub-millimeter SPECT was recently reported using a similar small animal system and a dedicated high-energy collimator (*15*). Additional improvements in dedicated high-resolution detectors will improve both sensitivity and resolution.

Future work will focus on the definition of optimal energy windows to maximize the recovery coefficient and sensitivity-versus-resolution tradeoffs of the SPECT system; and implementation of attenuation correction. Concurrent developments in collimator design, reconstruction parameters and detectors will enhance the scientific utility of these findings, and may have import to the direct translation of ^223^Ra in man. The use of noninvasive imaging as a means to predict and monitor therapeutic effect will enhance preclinical evaluation of altered dosing schedules and the role of combination treatments on ^223^Ra pharmacokinetics. We anticipate that these efforts will play a significant role towards improved patient treatment with ^223^Ra and other investigational alpha particle emitters.

## Acknowledgements

**Acknowledgements**

This work was supported in part by the Society of Nuclear Medicine and Molecular Imaging Junior Faculty Development Award (DSA), Steve Wynn Prostate Cancer Foundation-Young Investigator Award (DLJT) and David H. Koch PCF-YIA (DU); Knut and Allice Wallenberg Foundation (DU); Bertha Kamprad Foundation (DU); Patrick C. Walsh Foundation (DLJT) and the National Institutes of Health Johns Hopkins University Cancer Center Support Grant, P50-CA058236-19A1 (PI: Nelson), and RO1-CA201035 (DLJT).

## Methods

### Radium-223 and Characterization

Radium-223 dichloride was produced using a laboratory built Actinium-227 microgenerator, as previously detailed (*18*). Briefly, methanol/nitric acid was used to elute Radium-223 from parental ^227^Ac and ^227^Th. Using strong anion exchange chromatography, the purified material (solvated in 0.03 M citrate) was verified using a high purity germanium γ-spectrometer (Detective-EX-100, Ortec). For the representative spectra, a column fraction was placed on a platform at a distance of 2 cm from the detector and counted for 300 seconds. Spectra were initially analyzed in the Maestro software package (Ortec). A background spectra (in the absence of a radioactive sample) was also acquired for 360 sec with no detection of activity. Annotation of the spectra was provided from the Evaluated Nuclear Structure Data File hosted at the National Nuclear Data Center (*19*).

### Imaging System

The U-SPECT system (MILabs, Utrecht, Netherlands) is a dedicated preclinical imaging platform. It is comprised of three large panel (51 x 38 cm^2^) NaI detectors (9.5 mm thick crystals) with digital electronics providing full list mode acquisition, arranged in a triangular configuration. Subjects are placed in a multipinhole collimator via insertion on an automated and heated bed. The XUHS multi-pinhole tungsten collimator (54 pinholes of 2.0 mm diameter per pinhole, 48 mm tube diameter, MILabs) used in this study has a maximum resolution of 0.85 mm, and sensitivity of >13 kcps/MBq or 1.3% (*16*). The animal was placed on a heated imaging bed with integrated inhalation anesthesia which was translated through the center of the SPECT system for imaging a selected region of the animal (defined for these studies as the whole-body). Reconstructions consist of a pixel-based ordered subset expectation maximization algorithm, using the U-SPECT reconstruction software (*17,20*). Here, advanced resolution recovery based on a measured and interpolated position dependent point spread function compensates for distance dependent sensitivity and blur are applied within user selected energy windows (below). Attenuation correction was not applied. Scatter and background were corrected for with the triple-energy-window method (*21*).

### Phantom Studies

Line source phantoms each consisted of a capillary tubes (1.0 mm outer diameter, 0.5 mm inner diameter; Sutter Instrument Company) filled with ^223^RaCl_2_ in 0.03 M sodium citrate and stoppered with capillary tube sealant clay (Globe Scientific). Activity in each phantom was assessed using a CRC-127R well counter (Capintec Incorporated). Radium-223 was calibrated according to the guidelines provided by the supplier in response to the United States Nuclear Regulatory Commission (*22*). Specifically, the dial setting of the calibrator was determined to match a decay corrected National Institutes of Standards and Technology calibrated clinical dose of ^223^RaCl_2_ (Xofigo; Bayer HealthCare Pharmaceuticals). This empirically determined dial setting was #277.

The scattering phantom was constructed by casting a 1% (w/v) agar gel (Invitrogen) in a 10 mL syringe. When set, this formed a cylinder of 99% water with a diameter of 1.6 cm in which the capillary tube was embedded. The scattering phantom with the capillary tube was then centered in the field of view of the small animal SPECT system. Imaging of the phantom was performed using a single frame for 30 minutes. Post-acquisition reconstruction was performed with an energy window of 80-100 keV to capture the ^223^Ra γ-emissions at 81, 83.8, 94.3, 94.9 and 97.5 keV.

X-ray computed tomography (CT) was performed on a Sedecal SuperArgus R4 small animal PET/CT (Sedecal Systems). Reconstructed SPECT/CT fusions were generated using a three-dimensional data visualization suite registration package (Amira version 5.3.3, FEI Incorporated).

### Animal Studies

#### Radium-223 Imaging

The Institutional Animal Care and Use Committee of the Johns Hopkins University approved all procedures involving mice. C57Bl/6 mice were purchased from Charles River, all mice were male with weight between 22 and 27 g. As described above, animal images were acquired with the small animal USPECT system using a highly sensitive multipinhole collimator (the MILab XUHS collimator). Four frames were acquired of 22 minutes each, which were summed for reconstruction (using the energy windows described in Results). CT on the same animals was acquired as above, using the Sedecal SuperArgus R4.

#### Bone Scan

A representative mouse bone scan was acquired using a 44 mm inner-diameter, 1.0 mm diameter multipinhole collimator. The anesthetized mouse was administered 37 MBq ^99m^Tc-MDP (Cardinal Health) by retro-orbital injection. Imaging on the USPECT was performed 25 minutes after injection, using a single frame for 25 minutes.

#### Autoradiography

Whole body and hind limb specimens were flash frozen in optimal cutting temperature media (OCT, Sakura Fintec) and submerged in liquid nitrogen. Whole body autoradiography was performed using the Bright 8250 Cryostat (Bright Instruments) and sections of limbs were obtained using a modified cryotome (Leica 1800). En bloc images of sections (10 μm in thickness) were acquired, as previously described (*14*). Sections were exposed on phosphor screens and imaged using a desktop phosphor imaging scanner (Cyclone Phosphor Imager, Packard).

#### Histology

Reagents were purchased from ThermoFisher Scientific unless otherwise identified. Immunofluorescence labeling of fresh frozen sections was performed with γH2AX rabbit anti-mouse antibody (Ab-139, Sigma-Aldrich). Fresh frozen sections were bathed in room temperature 10% paraformaldehyde for 10 min, washed with phosphate buffered saline (PBS) and then blocked with goat serum for 1 h. The primary antibody was diluted to 1:250 in PBS and incubated overnight at 4° C. Alexa488-conjugated Goat anti-rabbit secondary was diluted 1:200 and applied to the PBS washed slides for 1 h at 4° C. Sections were counterstained with Hoechst 33342 for 5 minutes and washed in triplicate with PBS before mounting in an aqueous glycerol (30% v/v), and letting set overnight.

**Author Contributions**

Conceptualization, DLJT; Investigation, DSA, AR, RET, PAF, BWS, DLJT; Data Analysis, AR, BWS, DU, RCR, DLJT. Writing, All Authors; Supervision, BT, DLJT. The authors have no conflicts of interest relevant to this work.

## Supplemental Figures and Legends

**Supplemental Figure 1.**
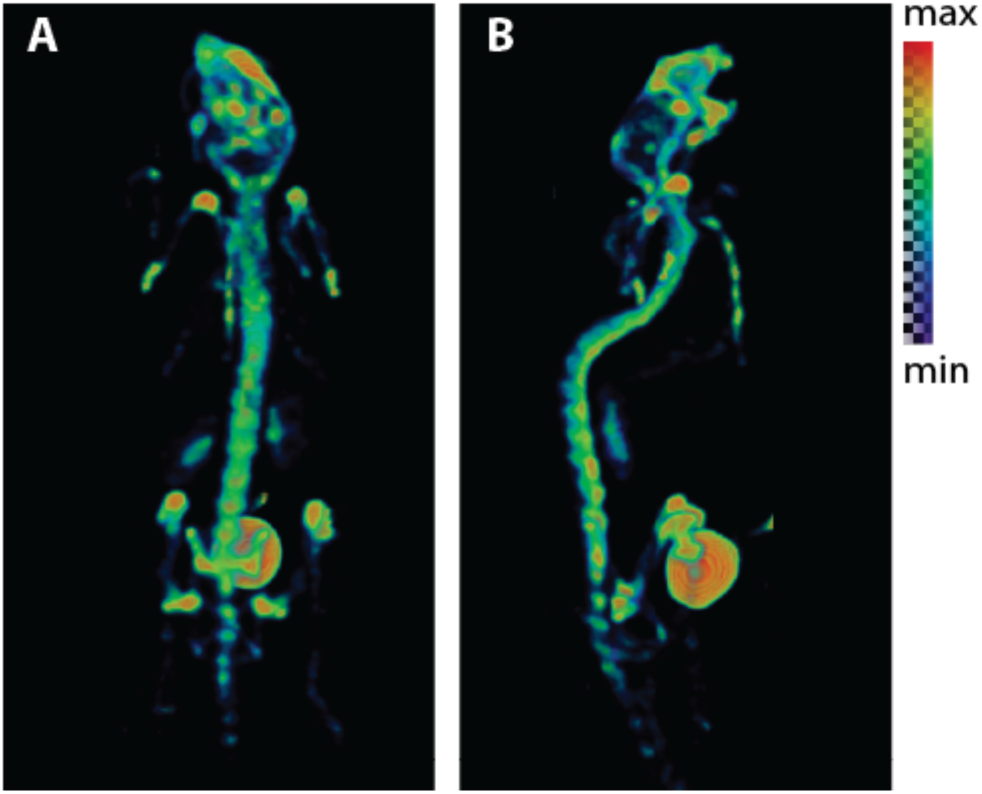
SPECT volumes of in A) dorsal and B) lateral views of a mouse 45 minutes after ^99m^Tc-MDP administration. The bone scan shows focal uptake throughout the skeleton, concentrated at the joints and skull, with significant signal accumulated in the bladder.

**Supplemental Figure 2.**
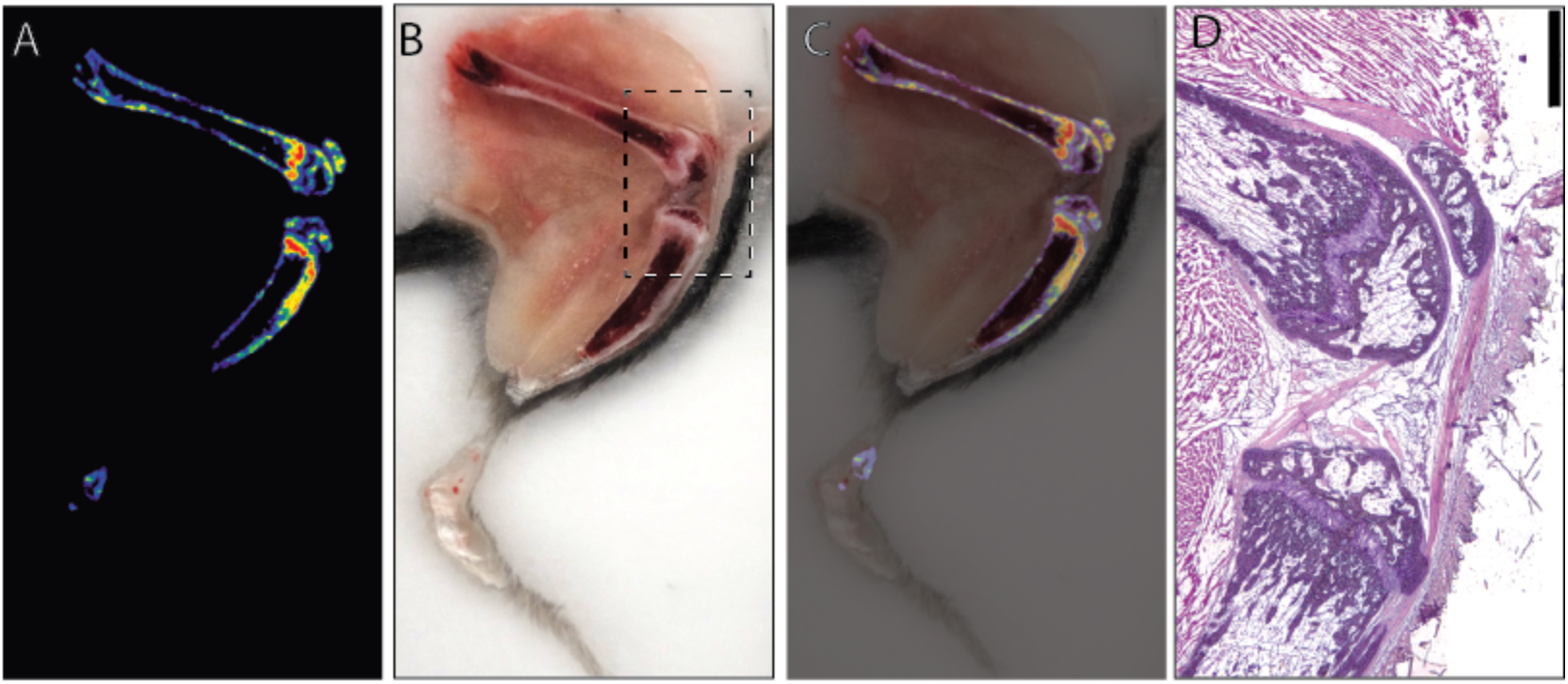
A) Autoradiography, B) cryo-sectioning and C) overlay of 48 h after administration of 0.55 *μ*Ci of ^223^Ra. D) Inset region of the cryo-section, stained by hematoxylin and eosin staining on the fresh undecalcified cryo-section (scale is 500 *μ*m). Uptake is localized to the unfused growth plates in the distal femur and proximal tibia, red arrows.

